# Impact of contamination on biology, biochemical, and histology of *Eobania vermiculata* and *Monacha obstructa* during different seasons in Ismailia, Egypt

**DOI:** 10.1101/2021.12.14.472665

**Authors:** Yassmeen S. M. Abd El mageed, Abd El Fattah Ali Ghobashy, Asma W. Al-Thomali, Maha F. M. Soliman, Amaal Mohammadein, Nahla S. El-Shenawy

**Affiliations:** Zoology Department, Faculty of Science, Suez Canal University, Ismailia, 41522, Egypt; Department of Biology, College of Sciences, Taif University, P.O. Box 11099, Taif 21944, Saudi Arabia

**Keywords:** land snails, morphometric, growth rate, proteins, lipid peroxidation, glutathione, histology

## Abstract

Land snails are found to be an appropriate sentinel organism, and the biomarkers chosen are effective for terrestrial heavy metal biomonitoring. The study aimed to compare the biological, biochemistry, and histology of two land snails in the Ismailia governorate, Egypt during different seasons. Random snails were collected from January 2015 to December 2015 from two sites in the Ismailia Governorate, on wet days during each season’s middle month. Soft tissues were taken from the dissected snails. It was noticed that most of the morphometric parameters measured shell height (ShH), last whorl width (LWW), maximum diameter (MaxD), aperture height (AH), and aperture width (AW) were higher in *Eobania vermiculata* (Sp. 1) than those measured in *Monacha obstructa* (Sp. 2), except for shell height measurement, which was the same in both species. The growth rate of Sp. 1 and Sp. 2 changed seasonally. In the more polluted areas with heavy metals, lipid peroxidation (LPO) was higher in snails and total protein content than in the snails collected from the less polluted areas for all seasons. However, the snails displayed lower levels of glutathione (GSH) as compared to snails at a less polluted site. GSH and LPO levels, on the other hand, have a negative relationship between them. Histopathological alterations in the digestive gland were more obvious in the general architecture of the digestive gland that had lost its tubular appearance. The excretory cells showed an increase in their excretory granules’ number and size while calcium cells decreased. Also, gonad follicles have lost their normal architecture with the degeneration of some stages of spermatogenesis and oogenesis. In conclusion, There was a strong correlation between GSH levels and total protein content in the same soft tissues. GSH and LPO levels, on the other hand, have a negative relationship. The overall results display the usefulness of *E. vermiculata* and *M. obstructa* land snails as bioindicator organisms and support the application of this ecotoxicological approach for evaluating the biologic impact of toxins. *E. vermiculata* is more abundant than *M. obstructa*. The density, morphometric, biochemical, and histology of *E. vermiculata* and *M. obstructa* were different at different seasons.

## 1. Introduction

Earthworms (Hankard et al., 2004), isopods (Nolde et al., 2006), and ground snails of the genera Cepaea, Helix (Regoli et al., 2006), as well as Xeropicta (Laguerre et al., 2009) and apple snails (Vega et al., 2012) have all been used to evaluate terrestrial pollution. The use of mollusks as emission sentinels is highly beneficial due to their widespread distribution, ease of sampling, stress resistance, and capacity to absorb a variety of contaminants (Anim et al., 2011). As a result of these findings, there is a growing interest in using land-adapted organisms as xenobiotic sentinels to forecast environmental protection.

The Mediterranean snail, *Eobania vermiculata*, has been designated as a sentinel species in this respect, as it has many advantages over other land snail species (Radwan et al., 2008; Toader-Williams and Golubkina, 2009; Itziou and Dimitriadis, 2011). Land snails are dispersed in Egypt in different governorates, which include Alexandria, Kafr El-Sheik, Behera, Damietta, Ismailia, Sharkia, Monofia, Gharbia, Minia, and Assiut (Shoieb, 2008; Ramzy, 2009; Eshra, 2013; Abdelgalil et al., 2018; Desoky, 2018). Many scholars have looked at ecological studies such as surveys, population dynamics and migration, daily life, and ground snail dispersal (Ramzy, 2009; El-Shenawy et al., 2012; Desoky et al., 2015).

This snail is widely spread in Egypt’s Ismailia region (Shoieb, 2008; Ramzy, 2009), implying that it may be used to estimate the pollution of the environment. According to Shoieb (2008), *Monacha obstructa* has been observed in the Ismailia Governorate, attacking a range of crops. The development of *M. obstructa* was spurred by human activity in the Middle East and the Arabian Peninsula, and it has since been recorded in a number of Middle Eastern nations, including Egypt, Saudi Arabia, Qatar, and Oman, as well as the Arabian Peninsula (Desoky et al., 2015).

In the warmth of the day, *E. vermiculata* climbs up trees, palms, bushes, and walls, and it can be found in all coastal areas, fields, plantations, vegetables, and vineyards (Radwan et al., 2008). The land snail is a good source of protein, and it is eaten by the vast majority of the population living in thick tropical forests and nearby areas, where snail farming is a major concern (Toader-Williams and Şara, 2010).

Due to their capacity to absorb trace metals in their tissues, land snails such as *H. aspersa* and *Arianta arbustorum* have historically been used to estimate urban or industrial waste (Regoli et al., 2006; Itziou and Dimitrids, 2009). *E. vermiculata* was found to be an appropriate sentinel organism, and the biomarkers chosen are effective for terrestrial heavy metal biomonitoring in Taif (El-Shenawy et al., 2012) as well as Ismailia (Ghobashy et al., 2021).

Non-enzymatic (GSH) and enzymatic [catalase (CAT), glutathione peroxidase (GPx), and glutathione-S-transferase (GST)] antioxidants were used to assess oxidative individual disturbances in the snails’ digestive glands. The loss of GSH in this species is connected to oxidative stress, and indicators for assessing contaminated terrestrial environments might be beneficial (Radwan et al., 2010). Protein loss is caused by catabolism of proteins in response to energy demands. Animals require a lot of energy to cope with stress, which can lead to protein catabolism. Furthermore, the decrease in protein levels might be linked to the creation of lipoproteins, which can be used to repair cell, tissue, and organ damage (Amiard and Amiard-Triquet, 2008).

Anthropogenic pollution can also be monitored using histopathology (El-Shenawy et al., 2007). Histological alterations of the digestive glands of snail mollusks’ tissues are responsive and sensitive to a wide range of contaminants because they play an important role in endocytosis of food substances, food absorption, storage, digestion, and excretion (Radwan et al., 2008).

Many investigations into the morphology and histology of the digestive gland of the land snail *E. vermiculata* have been conducted (Hamed et al., 2007; Radwan et al., 2008; Mohammadein et al., 2013). There are just a few studies that determine the number and kinds of cells that make up the digestive gland epithelium of terrestrial gastropods, despite the fact that the anatomy and physiology of digestive gland cells are well understood. This is mostly owing to the digestive gland epithelium’s multifunctional nature and the significant ultrastructural changes that occur during digestion (Radwan et al., 2008).

The morphological details of the shell provide information regarding their real body development (Roth and Mercer, 2000). Multivariate statistical studies of a collection of quantitative variables, such as the length, breadth, and height of a shell, were used in reported morphometrics. As the shell deteriorates, as it increases in size, shell material is added to the margins to keep its form, and at the top. The spiraling axis around which the aperture develops; it is frequently used as a shell indication expansion.

In our previous study, we showed that levels of essential (Cu, Mn, Zn, Co, Mo, Ca, and Mg) and non-essential (Cd, Pb, Al, and B) heavy metals in the soft tissues of two snail species (*E. vermiculata* and *M. obstructa*)collected from the Ring Road, Ismailia, Egypt, had the highest levels of heavy metals when compared to another site (El-Balah Road) (Ghobashy et al., 2021).

The present study aimed to compare *E. vermiculata* and *M. obstructa* by measuring the morphometric of the shell. Also, the determination of the biochemical markers related to oxidative stress (lipid peroxidation and glutathione) and energy reserves (total protein) for the *E. vermiculata* and *M. obstructa* at two sites through the four seasons was evaluated. Histological studies of ovotestis and digestive glands, as well as their histopathology, were considered during the research period.

## 2. Experimental procedures

### 2.1 Site of collection of snails and determination of population density

Ismailia, the north-eastern part of Egypt, is located approximately halfway between Port Said to the north and Suez to the south. Individuals of the herbivorous adult land snails, *E. vermiculata* (Müller) and *M. obstructa*, were collected from two sites in Ismailia, Egypt (supplemented material, 1). The first site (I) was El Dary Road, inside the Suez Canal University. This area is natural, whereas the area near the running cars is usually heavily polluted. The second site (II) was Al Balah Road, inside the old Suez Canal University. Also, We selected these two sites because the both species of snails are available. Four hundred individual snails were randomly collected from both sites each time, whereas the collection time was seasonal. Samples of snails, *E. vermiculata* and *M. obstructa*, were put in clean plastic jars and transported to the laboratory.

Adult snails, *E. vermiculata* and *M. obstructa*, were collected in the early morning from two sites in the Ismailia Governorate, Egypt, at approximately 5 am from January 2015 to December 2015. The quadrate count method has been used during the sampling procedure (0.5 m^2^X 0.5 m^2^) to measure the population density of each species in two studied sites during all four seasons (Dodd, 2011; Ghobashy et al., 2021). The snails were collected three times from each site seasonally and transferred to the Zoology Department, Faculty of Science, Suez Canal University, Ismailia, in plastic bags. The snails were identified according to the key given by Radwan et al. (2008). The individual species’ absolute population density was calculated as follows:

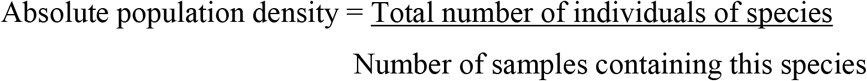

#### Ethics approval

Zoology Department, Faculty of Science, Suez Canal University was approved this research.

### 2.2 Morphometric measurements

A total of 100 snails were used from each species seasonally. Five shell parameters were taken in either the axial view [shell height (ShH), last whorl width (LWW), and maximum diameter (MaxD)] or the apertural view [aperture height (AH) from the point of adhesion of the aperture to the body whorl and aperture width (AW)] (Figs. 2A) to describe quantitatively the size and shape using a Vernier caliper (to the nearest centimeter). Only shells with a reflected lip were used because this signals the snail’s maturity as well as the end of its growth cycle (Madec et al., 2003).

### 2.3 Tissue preparation

The animals were washed with distilled water. The shells of non-anesthetized tested snails were removed by continuously cutting the whorls starting at the aperture opening using strong scissors, and the broken fragments of the shell were carefully removed. Soft tissues were rapidly dissected out (Itziou and Dimitriadis, 2009) from one hundred chosen snails at both sites. At each time point, 80 animals were stored at -20 °C until they were used in the biochemical analysis. For the histological study, snails (n = 20) were rapidly excised and immediately dropped into suitable fixatives.

### 2.4 Biochemical determination

Determination of lipid peroxidation (LPO) in the tissue was used as an oxidative indicator parameter. The evaluation was based on the measurement of thiobarbituric acid reactive product malondialdehyde (MDA) in the acidic medium at a temperature of 95 °C for 30 min, according to Draper and Hadley (1991). The content of glutathione (GSH) in snail tissues was estimated immediately using the Beutler et al. (1963)method. Proteins were measured using the total protein kit according to the Biuret method (Bio-diagnostic, 29 Tahreer St., Dokki, Giza, Egypt). The creation of colored complexes between peptide bonds and cupric ions in an alkaline medium is the basis for this process.

### 2.5 Histological study

For histological studies, the gonad and digestive gland of snails from different stages were quickly excised and immersed in formalin solutions (10%) for 48 hr. The thin sections of 4μm were obtained by semiautomatic microtome (Kiernan, 1990). The slides were stained with eosin and hematoxylin and observed under a light microscope.

### 2.6 Statistical analysis

The statistical analysis was performed using SPSS statistics, version 17.0. The mean values obtained in the different groups were compared using the two way ANOVA test combined with the post-hoc Turkey test to determine the significant difference between species at different sites. The simple positive/negative correlation analysis (Pearson’s test) was applied to examine the relationship between the biochemical parameters and the concentrations of heavy metals as well as the rate of growth and type of food.

## 3. Results

### 3.1 Population density

Table 1depicts a study of the population density of land snail species at both sites in the Ismailia Governorate. The two land snails, *E. vermiculata* (supplemented material, 2a) and *M. obstructa* (supplemented material, 2b), were found in the two sites described earlier throughout the year. The brown garden snail, *E. vermiculata*, was found to be the most common species at the two sites investigated in a survey with a relatively high number during the four seasons. Generally, it was concluded that during the spring season, the population of *E. vermiculata* and *M. obstructa* increased as compared to the population during summer and autumn. The lowest population density was observed during the winter season.

**Table 1:** **Population density of *E. vermiculata* and *M. obstructa* in two study sites in Ismailia governorate during four seasons from January 2015 to December 2015.**

| Seasons | Mean number of snails per sample |  |  |  |
| --- | --- | --- | --- | --- |
|  | Site I |  | Site II |  |
|  | Sp. 1 | Sp. 2 | Sp. 1 | Sp. 2 |
| **Winter** | 100<sup>d, e, f</sup> | 16<sup>b, d, e, f</sup> | 104<sup>d, e, f</sup> | 18<sup>b, d, f</sup> |
| **Spring** | 160<sup>c, e, f</sup> | 40<sup>b, c, e, f</sup> | 154<sup>a, c, e, f</sup> | 36<sup>b, c, e, f</sup> |
| **Summer** | 130<sup>c, d, f</sup> | 24<sup>b, c, d, f</sup> | 130<sup>c, d, f</sup> | 22<sup>b, d, f</sup> |
| **Autumn** | 140<sup>c, d, e</sup> | 30<sup>b, c, d, e</sup> | 136<sup>c, d, f</sup> | 28<sup>b, c, d</sup> |

Data presented as the mean number of snails (n=3) where site I: New Suez Canal University (Ring road), site II: Old Suez Canal University (El Balah Road), sp. 1: *E. vermiculata*, sp. 2: *M. obstructa*, <sup>a</sup> Significant difference as compared to the site I, <sup>b</sup> significance difference as compared to sp. 1, <sup>c</sup> significance difference as compared to the winter season, <sup>d</sup> significant difference as compared to the spring season, <sup>e</sup> significant difference as compared to the summer season and significant difference as compared to the autumn season (P<0.05).

### 3.2 Morphometric measurements and growth rate

The growth rate was determined using different shell parameters. Table 2 presented the mean values of aperture height (AH), aperture width (AW), last whorl width (LWW), maximum diameter (MaxD), and shell height (Sh.H). It was noticed that most of the morphometric parameters measured (AH, AW, MaxD, and LWW) were higher in Sp.1 (*E. vermiculata*) than those measured in Sp. 2 (*M. obstructa*), except for shell height measurement, which was the same in both species.

**Table 2:** **Morphometric measurements of *E. vermiculata* and *M. obstructa* in two study sites of Ismailia governorate during four seasons from January 2015 to December 2015.**

| Morphometric parameters |  |  |  |  |  |  |  |
| --- | --- | --- | --- | --- | --- | --- | --- |
| Season | Site | Species | AW | AH | MaxD | LWW | ShH |
| Winter | Site I | Sp.1 | 1.36 ± 0.02 <sup>d, f</sup> | 1.38 ± 0.03 | 2.17 ± 0.02 <sup>d, e, f</sup> | 1.14 ± 0.03 <sup>d, e</sup> | 1.41 ± 0.03 |
|  |  | Sp.2 | 1.42 ± 0.03 <sup>e</sup> | 1.40 ± 0.03 <sup>e</sup> | 2.09 ± 0.03 <sup>d, f</sup> | 1.21 ± 0.04 <sup>e</sup> | 1.41 ± 0.04 <sup>e</sup> |
|  | Site II | Sp.1 | 1.38 ± 0.02 <sup>f</sup> | 1.34 ± 0.02 <sup>f</sup> | 2.16 ± 0.03 <sup>d, e, f</sup> | 1.16 ± 0.02 <sup>e</sup> | 1.42 ± 0.02 |
|  |  | Sp.2 | 1.36 ± 0.02 | 1.38 ± 0.02 | 2.09 ± 0.02 <sup>d</sup> | 1.18 ± 0.03 | 1.40 ± 0.01 <sup>e</sup> |
| Spring | Site I | Sp.1 | 1.55 ± 0.01 <sup>c</sup> | 1.46 ± 0.02 | 2.49 ± 0.02 <sup>c</sup> | 1.37 ± 0.02 | 1.51 ± 0.02 |
|  |  | Sp.2 | 1.50 ± 0.02 | 1.42 ± 0.02 <sup>e</sup> | 2.44 ± 0.02 <sup>c, e</sup> | 1.31 ± 0.02 <sup>e</sup> | 1.51 ± 0.02 <sup>e</sup> |
|  | Site II | Sp.1 | 1.50 ± 0.02 <sup>c</sup> | 1.45 ± 0.01 | 2.46 ± 0.02 <sup>c</sup> | 1.16 ± 0.02 | 1.46 ± 0.02 |
|  |  | Sp.2 | 1.50 ± 0.02 <sup>c</sup> | 1.43 ± 0.02 <sup>e</sup> | 2.51 ± 0.03 <sup>c, e, f</sup> | 1.18 ± 0.03 | 1.49 ± 0.02 <sup>e</sup> |
| Summer | Site I | Sp.1 | 1.46 ± 0.02 | 1.41 ± 0.03 | 2.41 ± 0.02 <sup>c</sup> | 1.30 ± 0.02 <sup>c</sup> | 1.44 ± 0.02 |
|  |  | Sp.2 | 1.29 ± 0.06 <sup>b, c, d</sup> | 1.26 ± 0.06 <sup>b, c, d</sup> | 2.15 ± 0.11 <sup>b, d, f</sup> | 1.06 ± 0.07 <sup>b, c, d, f</sup> | 1.27 ± 0.06 <sup>b, c, f</sup> |
|  | Site II | Sp.1 | 1.46 ± 0.03 | 1.41 ± 0.03 | 2.45 ± 0.03 <sup>c</sup> | 1.36 ± 0.07 <sup>c, d</sup> | 1.44 ± 0.02 |
|  |  | Sp.2 | 1.30 ± 0.05 <sup>b, d</sup> | 1.26 ± 0.05 <sup>b, d</sup> | 2.18 ± 0.08 <sup>b, d</sup> | 1.08 ± 0.05 <sup>b</sup> | 1.26 ± 0.04 <sup>b, c</sup> |
| Autumn | Site I | Sp.1 | 1.56 ± 0.02 <sup>c</sup> | 1.46 ± 0.03 | 2.51 ± 0.03 <sup>c</sup> | 1.25 ± 0.02 | 1.47 ± 0.02 |
|  |  | Sp.2 | 1.44 ± 0.04 <sup>e</sup> | 1.39 ± 0.04 | 2.39 ± 0.06 <sup>b, c</sup> | 1.22 ± 0.04 <sup>e</sup> | 1.44 ± 0.04 <sup>e</sup> |
|  | Site II | Sp.1 | 1.58 ± 0.02 <sup>c</sup> | 1.52 ± 0.02 | 2.58 ± 0.03 <sup>c</sup> | 1.23 ± 0.01 | 1.51 ± 0.02 |
|  |  | Sp.2 | 1.39 ± 0.05 <sup>b</sup> | 1.31 ± 0.04 <sup>b</sup> | 2.23 ± 0.08 <sup>b</sup> | 1.15 ± 0.04 | 1.33 ± 0.04 <sup>b</sup> |

Data presented as mean ± S.E. (n=20) where: aperture height (AH), aperture width (AW), last whorl width (LWW), maximum diameter (MaxD), and shell height (Sh.H), the site I: Ring road, site II: El Balah Road, sp. 1: *E. vermiculata*, sp. 2: *M. obstructa*. <sup>a</sup> Significant difference as compared to the site I, <sup>b</sup> significance difference as compared to sp. 1, <sup>c</sup> significance difference as compared to the winter season, <sup>d</sup> significant difference as compared to the spring season, <sup>e</sup> significant difference as compared to the summer season and <sup>f</sup> significant difference as compared to the autumn season (P<0.05).

During the autumn season, the MaxD of Sp.1 was 2.58 cm, the maximum AW reached 1.58 cm, and the maximum AH was 1.52 cm. However, the greatest LWW was1.37 cm in the spring season, while, the highest Sh.H value was 1.51 cm in both the spring and autumn seasons. During the spring season, Sp. 2 attained its maximum size in all morphometric parameters. There was no significant difference between the two sites in terms of morphometric characteristics (P > 0.05). All morphometric parameters assessed for both species were likely to be strongly linked.

### 3.3 Biochemical of soft tissues

Figs. 1, 2 and 3 demonstrate the amounts of GSH, total protein, and lipid peroxidation in the soft tissue of two species of land snails throughout four seasons at two distinct locations in Ismailia, respectively. GSH levels did not differ significantly between two species on the same site or between two locations. The levels of GSH, on the other hand, were significantly different (P ≤ 0.01) between the four seasons. Throughout the summer, GSH reached its highest concentration for two species at the two sites. During the autumn, the GSH concentration was reduced significantly at site I for the two species as compared to site II (Fig. 1).

**Fig. 1:**
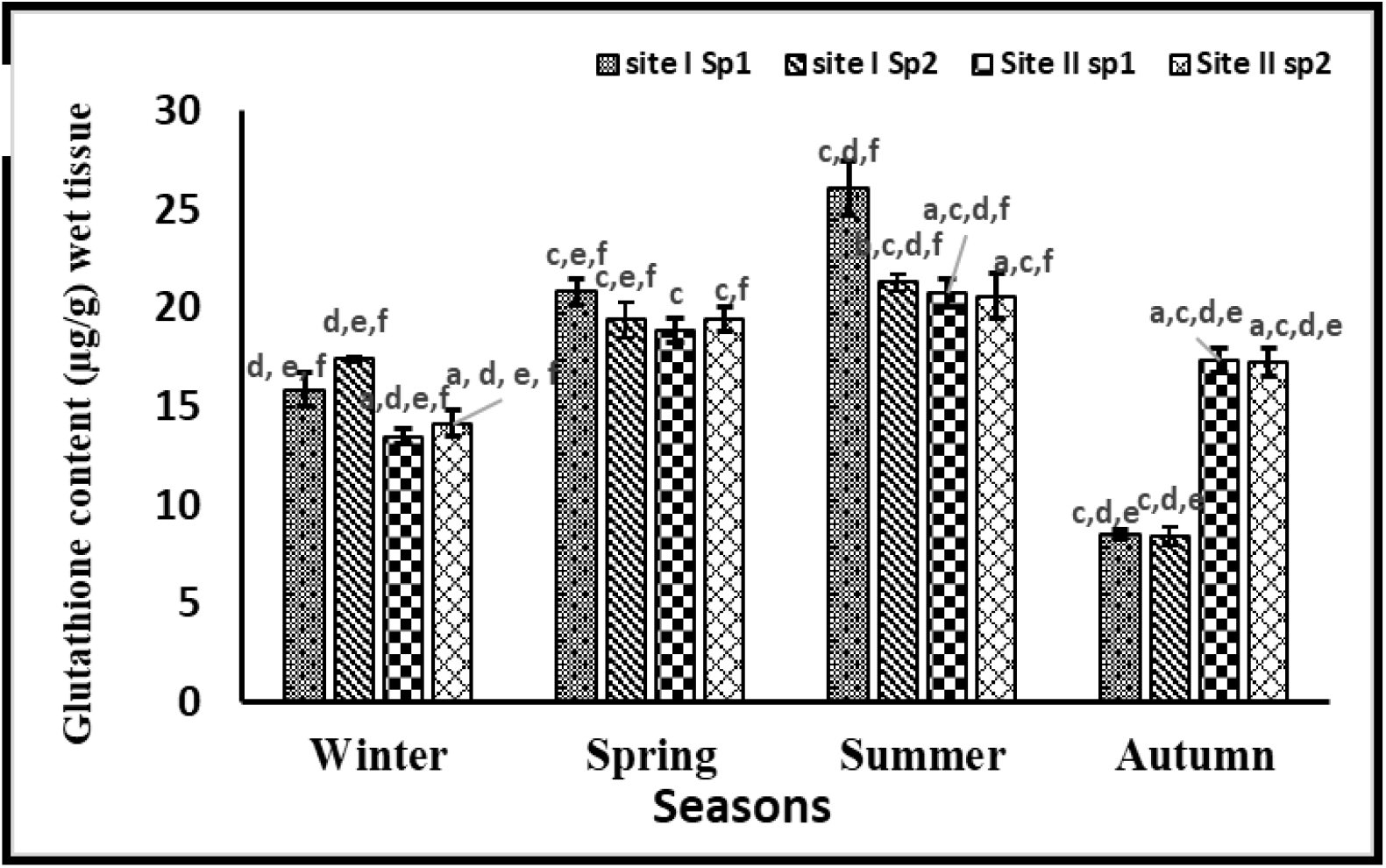
Glutathione content (μg/g) in the soft tissues of two species of land snails at two sites. The data presented as mean ± S.E (n= 5 samples). Site I: Ring road, Site II: El Balah road. ^a^Significant difference as compared to the site I, ^b^significance difference as compared to Sp. 1, ^c^significant difference as compared to the winter season, ^d^significant difference as compared to the spring season, ^e^significance difference as compared to the summer season, and ^f^significant difference as compared to the autumn season (P<0.05).

The total protein levels of the different species did not distinguish substantially across the two locations or during the four seasons. Unless total protein levels at sites I and II somewhat rise in the winter and spring seasons, respectively (Fig. 2).

**Fig. 2:**
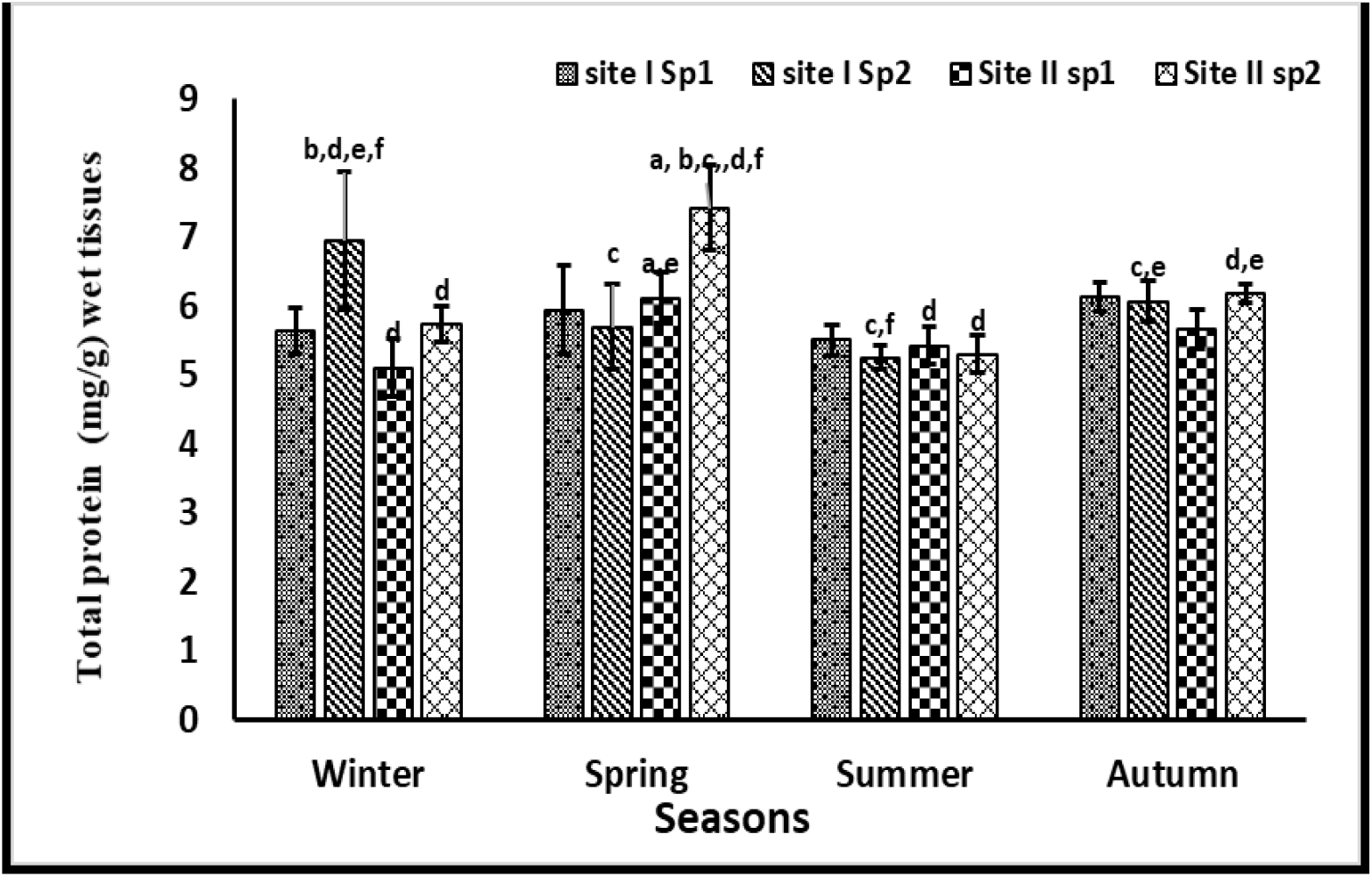
Total protein level (mg/g) in the soft tissues of two species of land snails in two sites. The data presented as mean ± S.E (n= 5samples). Site I: Ring road, Site II: El Balah road. ^a^Significant difference as compared to the site I, ^b^significance difference as compared to Sp. 1, ^c^significance difference as compared to the winter season,^d^significant difference as compared to the spring season, ^e^significant difference as compared to the summer season, and ^f^significant difference as compared to the autumn season (P<0.05).

LPO levels were found to be somewhat higher in the winter for two species at site-I than in the spring and summer. However, LPO levels of both species were found to be somewhat greater in the autumn at site II (Fig. 3). There was a strong correlation between GSH levels and total protein content in the same soft tissues. GSH and LPO levels, on the other hand, have a negative relationship.

**Fig. 3:**
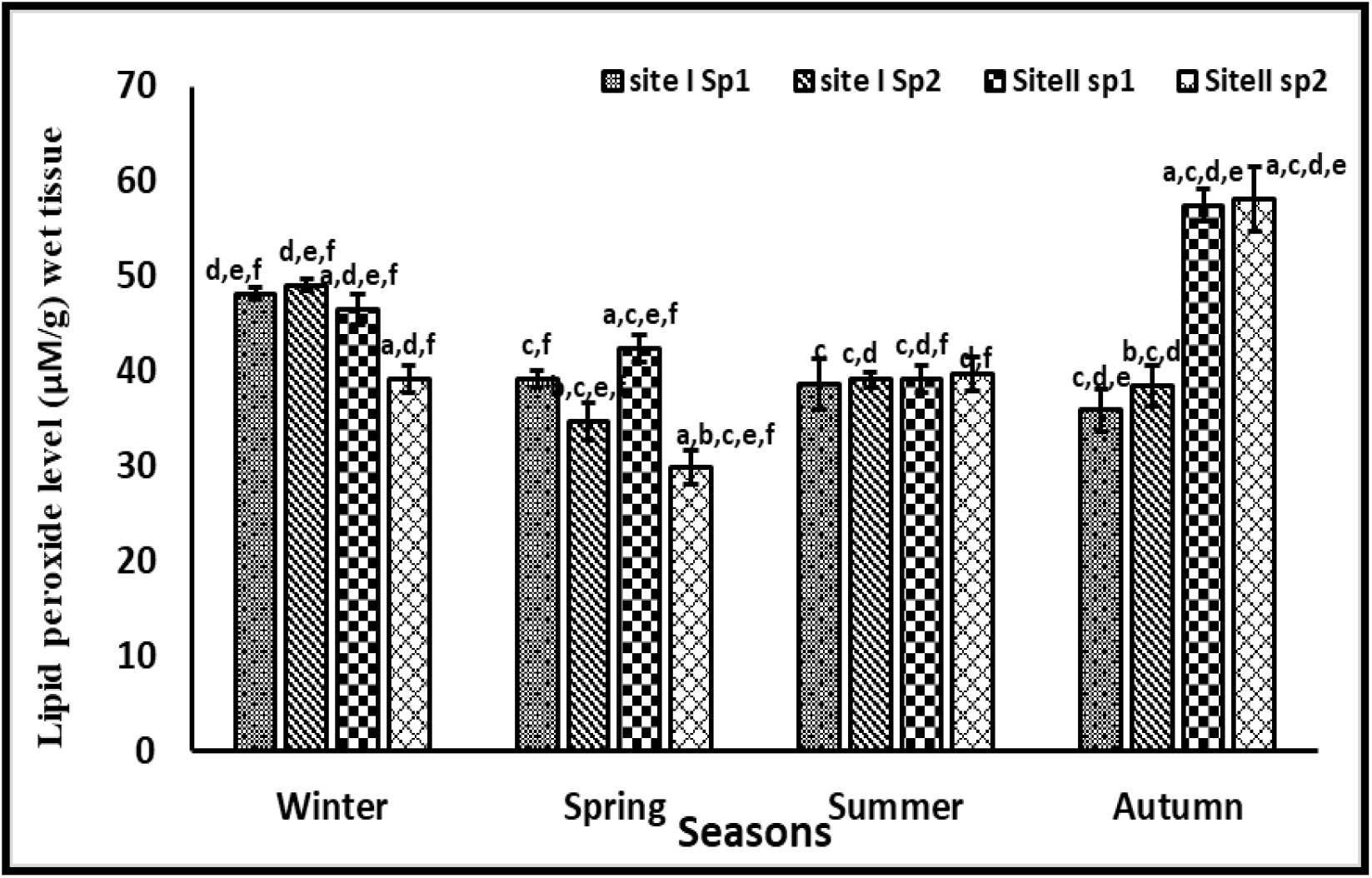
Lipid peroxide level (mg/g) in the soft tissues of two species of land snails in two sites. The data presented as mean ± S.E (n= 5samples). Site I: Ring Road, Site II: El Balah Road. ^a^Significant difference as compared to the site I, ^b^significant difference as compared to Sp. 1, ^c^significance difference as compared to the winter season, ^d^significant difference as compared to the spring season, ^e^significant difference as compared to the summer season, and ^f^significant difference as compared to the autumn season (P<0.05).

### 3.4 Histological and histopathological of the gonad and digestive gland

A hermaphroditic gland (ovotestis) is found in the posterior part of the digestive gland near the apex of *E. vermiculata* and *M. obstructa’s* reproductive system. The ovotestis is made up of a number of follicles or acini implanted in the digestive gland. Each acinus comprises of primary oogonia and spermatogonia in the early stages. Oogonia were large and rounded compared to spermatogonia, which were small in size (Fig. 4). Gonad follicles lost their normal architecture with degeneration of several phases of spermatogenesis and oogenesis, according to histopathological abnormalities in samples (Fig. 5a). Due to significant destruction of germ cells and additional degenerative consequences, acini seemed to be empty and replaced by brown granules (Fig. 5b).

**Fig. 4:**
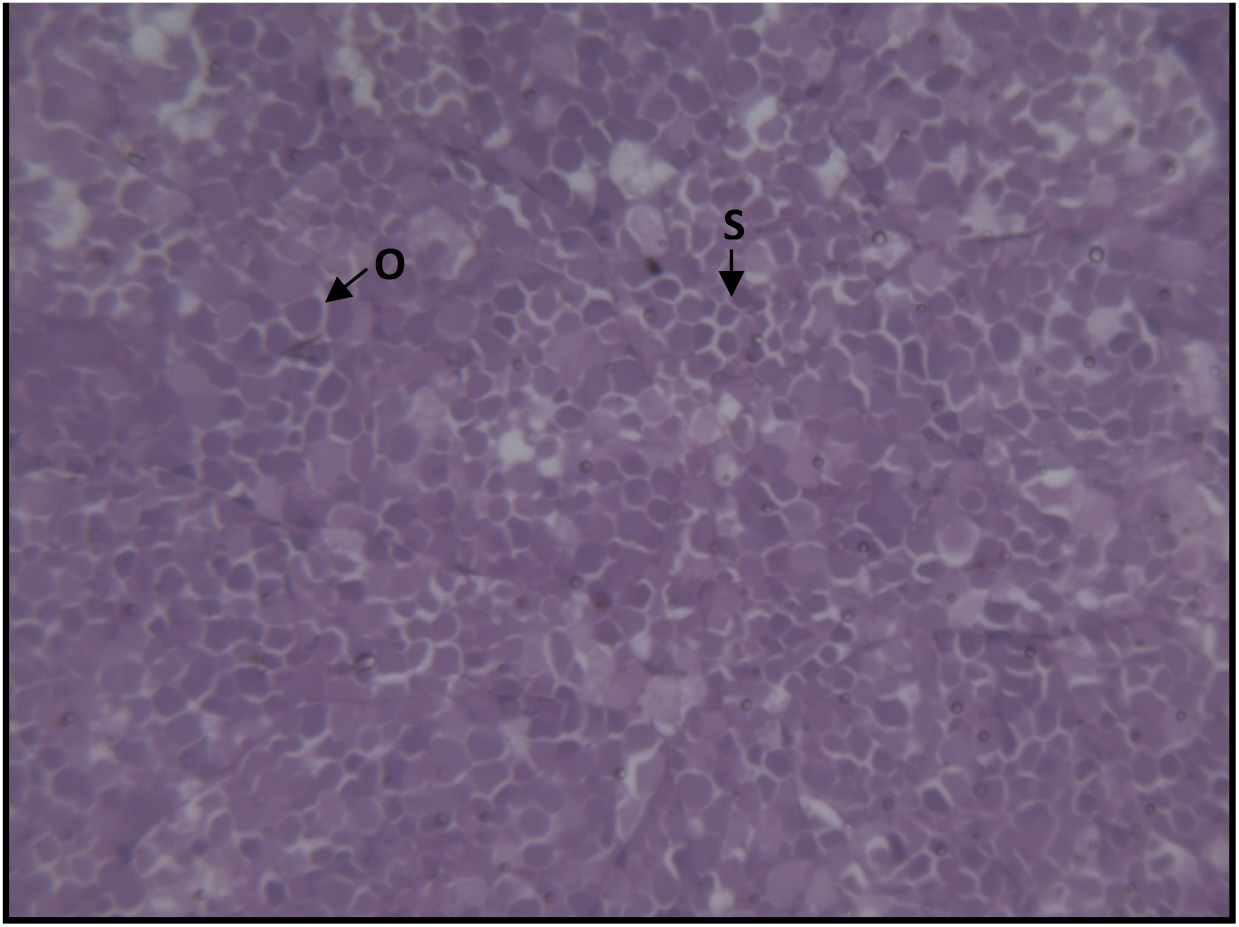
Transverse sections in ovotestis of two species of land snails, showing early stages of maturation. O: oogonia and S: spermatogonia (X=200).

**Fig. 5:**
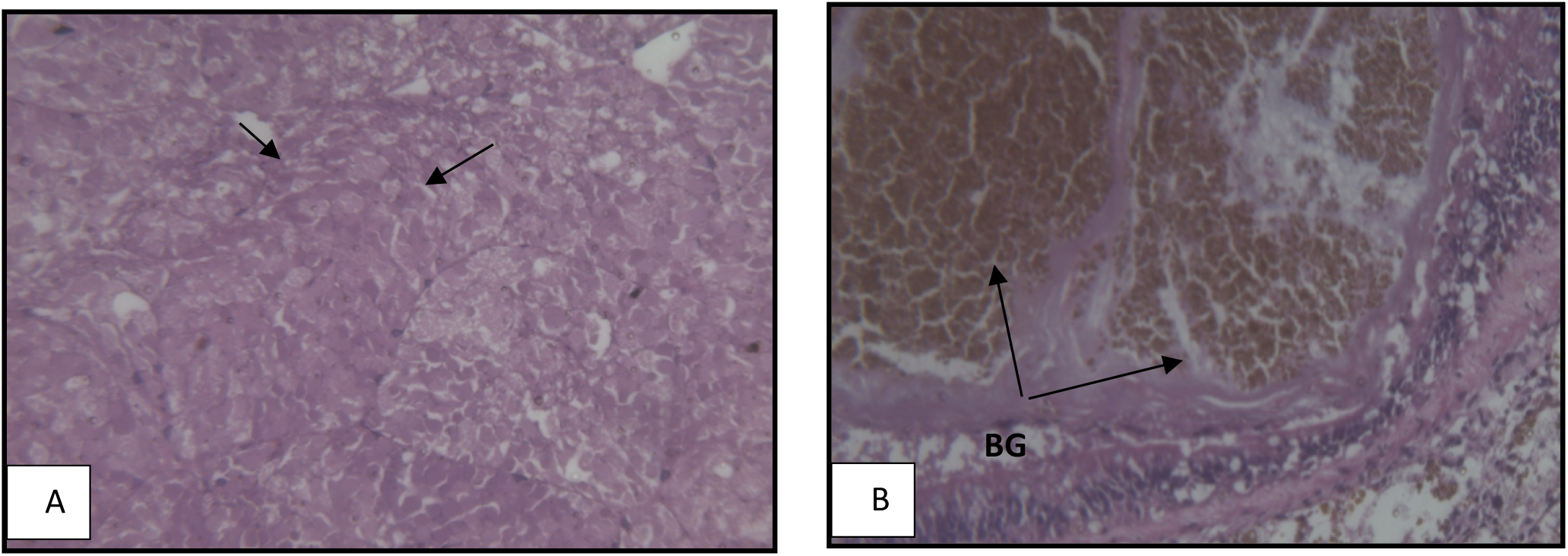
Transverse sections in ovotestis of two species of land snails. A: degeneration of spermatogonia and oogonia (arrows). B: tissue necrosis (arrow) and a large number of brown granules (BG) (X=200).

The digestive glands of *E. vermiculata* and *M. obstructa* are quite big, dark brown, and take up a significant portion of the snail’s visceral hump. The tubules of this gland are highly branching and blindly terminating tubules that are linked together by connective tissue (Fig. 6a). Each tubule of the digestive gland is surrounded by a thin layer of circular muscle fiber (Fig. 6b). When these cells were investigated, three types of cells were found. The cells of digestion, calcium, and excretion. Digestive cells are columnar cells with numerous vacuoles that accommodate yellowish brown granules and rounded or oval nuclei at the base. Excretory cells are spherical cells with a tiny nucleus and big dark brown excretory granules. Calcium cells are pyramidal in shape, with spherical calcium spherules (looking as luminous bodies) piled in the center and globular nuclei.

**Fig 6:**
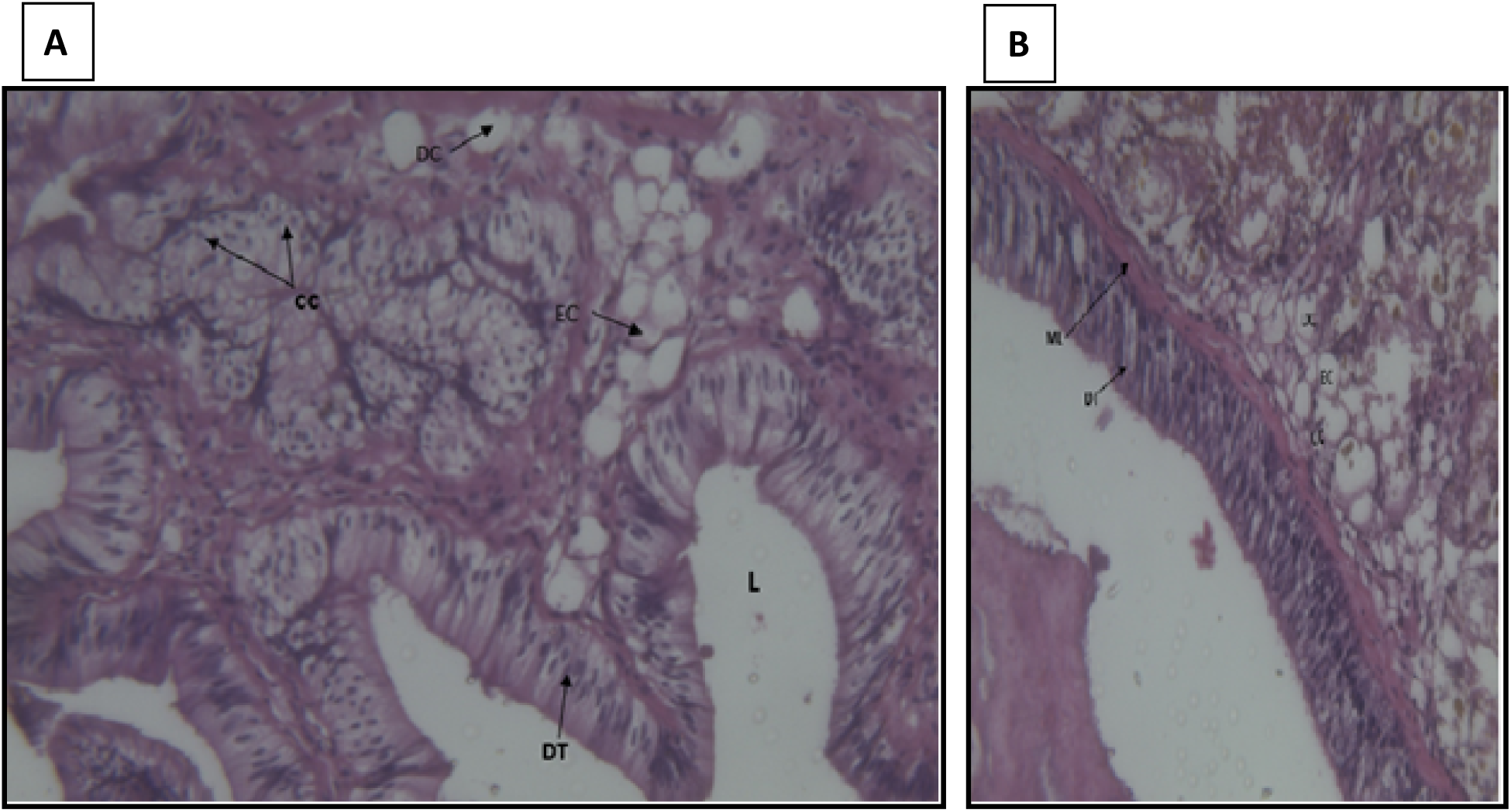
Transverse sections in the digestive gland of two species of land snails. **A:**the highly branched tubules of the digestive gland (DT) and three types of cells, digestive cell (DC), excretory cell (EC), and calcium cell (CC). Lumen (L) **B: a**thin layer of circular muscle fiber (ML) (X= 200).

The tubules of the digestive gland lose their typical morphology in polluted areas. On the other hand, the number and size of excretory granules in excretory cells rose, whereas calcium cells dropped and digestion cells were badly damaged, becoming considerably vacuolated (Fig. 7A). Calcium cells were packed with enlarged calcium spherules and exhibited pyknotic nuclei (PN). The cytoplasm of many calcium cells had been replaced by large vacuoles containing darkly pigmented granules. Excretory cells had more excretory granules, as well as cellular debris and necrotic regions (NA). The majority of tubules had been destroyed (Fig. 7B). The cytoplasm of digestive cells tended to be almost devoid of cytoplasmic granules, as well as the number of calcium cells and the size of calcium spherules were significantly reduced. Also, calcium cells contain an increased deposition of calcified brown granules (BG) with vacuolated cytoplasm (V) and karyolitic nuclei (KN). Hemocytic infiltration (HI) was frequently observed (Fig. 7C).

**Fig. 7:**
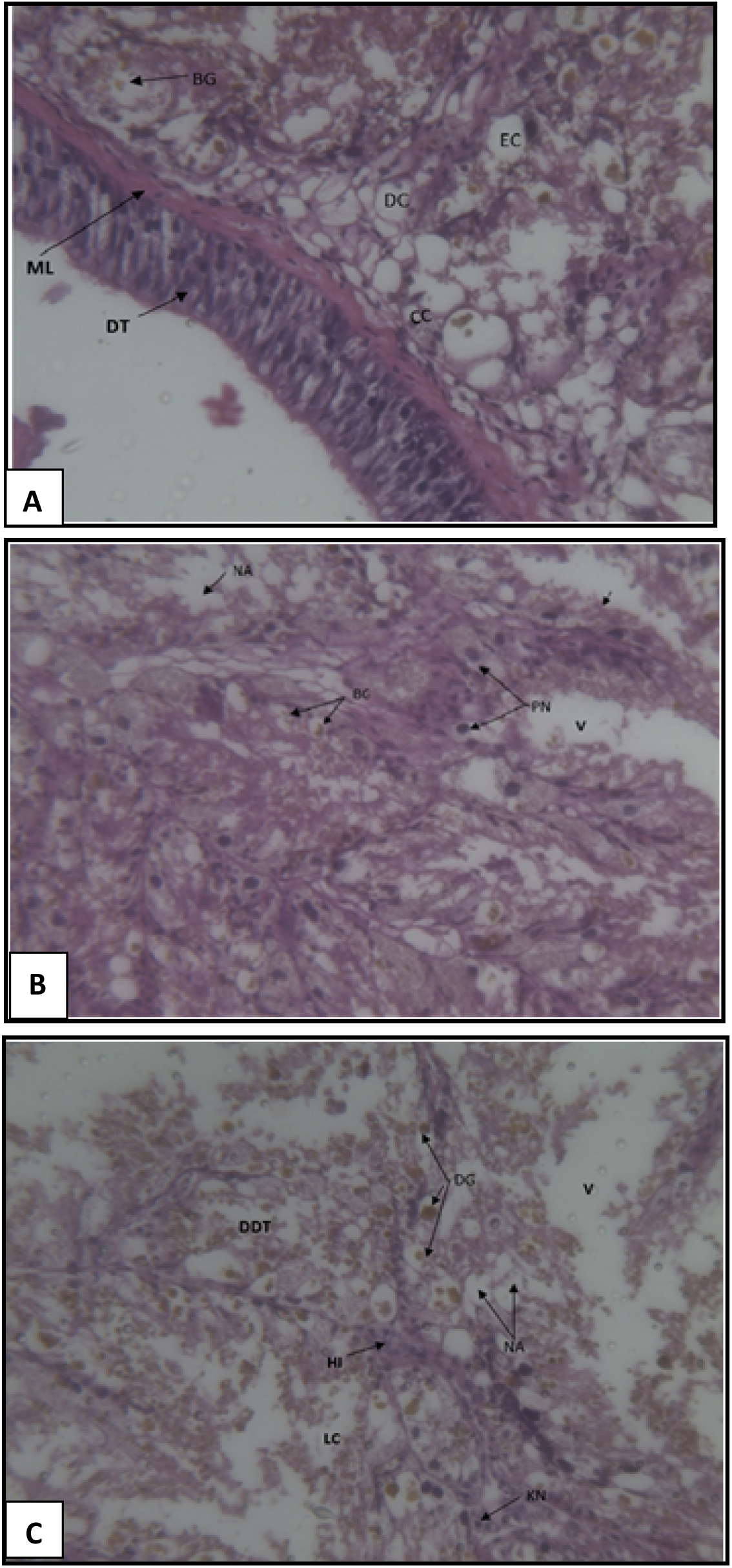
Transverse sections in the digestive gland of two species of land snails. **A:**digestive gland showing the tubules of the digestive gland lost their normal appearance and the excretory cells increased (EC) while calcium cell (CC) decreased. **B:**digestive cells loaded with brownish granules (BG) while the excretory cells are occupied by large vacuoles (V), the basement membrane of tubules appeared ruptured (arrow) and exhibited pyknotic nuclei (PN) and necrotic areas (NA). **C:**calcium cells contained an increased deposition of calcified brown granules (BG) with vacuolated cytoplasm (V) and karyolitic nuclei (KN). Hemocytic infiltration (HI) was frequently observed (X= 200).

## 4. DISCUSSION

The results revealed that both *E. vermiculata* and *M. obstructa* were found all over the year at the two sites mentioned in the Ismailia governorate. Land snails were previously distributed in the North Delta, where suitable climatic conditions (temperature-humidity-plant cover) existed, and with climate change, they have begun to spread to governorates in southern Egypt (Giza, BeniSuef, Minia, and Assiut). Desoky (2018)found that *E. vermiculata* and *M. obstructa* were recorded in Sohage for the second time. More widespread and maybe this pest has moved to these governorates with transportation. Passengers from places spread to these new places and adapted to the new region. *M. obstructa* has also been found in the governorates of Ismailia and Kafr El-Sheikh (Gabr et al., 2006; Shoieb, 2008). *E. vermiculata, M. obstructa, O. elegans, Vallonia pulechella, T. pisana, Vitrea pygmaea, Helicodiscus singleyanus inermis, Pupoides coenopictus*, and *Cecilioides acicula* were among the nine land snail species studied by Ramzy (2009)in the Assiut Governorate. Abo-El-Naser (2013)found that four terrestrial snails include three land snails (*E. vermiculata, M. obstructa*, and *oxyloma elegans*) and slugs (*Limax flavus*) in Assiut Governorate.

Mohamed (2015)reported that ten land snail species, *E. vermiculata, Theba pisana, Helicella vestalis, Cochlicella acuta, M. cartusiana, M. obstructa, Succinea putris, Succinea oblonga, Achatinidae*, and *Oxychilus alliarius* were recorded on different vegetation, vegetable, fruit, and ornamental plants in the North East of the Delta. All previous studies on the population density of both species have shown that *E. vermiculata* and *M. obstructa* snails are the most dominant snail species in the Egyptian governorates attacking various plantations (Eshra, 2013), so they were used in the current study.

The *vermiculata* land snail was the predominant species at the two studied sites in the Ismailia governorate. These findings are consistent with those of Eshra (2013), who investigated the spread of terrestrial snails in fruit orchards and ornamental plants in Egypt’s governorates of Alexandria and El-Beheira. *Theba pisana* and *E. vermiculata* were the most abundant in all regions, while *M. obstructa* snails were found at relatively low population densities. Shahawy (2019)also reported that *E. vermiculata* was the predominant species in Kafr El-Sheikh governorate on ornamental plants and was recorded on all surveyed plants in relatively high numbers. However, the present data disagrees with that reported by Rady et al. (2014), who stated that the *M. cartusiana* land snail was the predominant species in the fields of vegetable crops in the Ismailia and Sharkia governorates. Also, Rady et al. (2019)recorded that *M. cartusiana* was the predominant species in Qalubia and Sharkia governorates. Land snail population density, on the other hand, varies from one host plant to the next.

According to the findings of this study, the population of *E. vermiculata* increased during the spring months compared to the population during the summer and autumn months, when the weather conditions are favorable for snail activity. These findings corroborate Shahawy’s findings (2019) and Awad (2014), who found that the population fluctuations of land snails varied according to crop, temperature, relative humidity, and season. Snails are more active in the spring and autumn, and their numbers are lower than in the winter. The activity of land snails is also influenced by moisture.

The mucus of land snails consists of 98% water. As a result, water deficiency inhibits high-temperature activity. Light influences the activity of land snails; they remain in their hides during the day and only after dusk to search for food. Snails moved to foliage as the light intensity decreased due to a drop in temperature below 21 °C and an increase in humidity due to dewfall at dusk, while in the morning and during the day they returned to the upper soil or between the earth clods where it is cold and shady (Daoud, 2004; Ramzy, 2009). In the current study, *E. vermiculata* reached its maximum size (2.58 cm MaxD) in the autumn season. These observations are consistent with those of Georgiev et al. (2005), who found that *Heliccella obvia* reached its maximum size in October and November. Al-Khafaji et al. (2016)recorded that *E. vermiculata* reached a maximum shell diameter of 33 mm in Basrah, Iraq, in contrast with current data, which is much smaller. Climate conditions such as the temperature gradient co-varying with different seasons (Welter-Schultes, 2000) and humidity were blamed for the differences in shell size (Giokas, 2008), as well as arising from selection in different environments and exposure to different environmental factors (Welter-Schultes, 2000).

In the current study, there was no significant difference in morphometric parameters of two species of land snails between the studied sites. This could be due to the close distance between the two studied sites and since they are exposed to the same environmental conditions. The shell morphology is similar to that described by Georgiev et al. (2005), who recorded that it was almost spherical. Creamy-white on the underside, with a shiny earlobe-shaped aperture. Two honey-colored bands and two creamy-white, as well as striated bands on the ends. The shell is usually creamy white to pale yellow. Up to five brown spiral bands can be found, which are often fused. It’s possible to find specimens that are almost maroon brown. Usually, the bands are broken up by the white axial fire, resulting in an overlapping pattern of light and dark colors on the shell’s back. As a result, further research is required to determine the relationship between the shell shape and morphology of the studied organisms and environmental factors.

In the present data, significantly elevated levels of LPO in the soft tissue of *E. vermiculata* snails collected from the Ring Road in the winter season indicate that some cell damage might have occurred. This result may be due to the higher concentration of most essential heavy metals and Pb in the winter season of the study locations (Ghobashy et al, 2021). These findings were consistent with the current evidence, which showed that elevated Pb levels increased LPO levels in the digestive gland of polluted-site samples. Increasing LPO levels during exposure to metals have been recorded in several organisms (Radwan et al., 2010; El-Shenawy et al., 2012). Several studies have looked into the toxic effects of Pb on membrane components and have discovered a connection between these effects and oxidative damage caused by Pb. When Pb was incubated with various polyunsaturated fatty acids, a marked enhancement in MDA concentration was observed. As the amount of double bonds in fatty acids increased, the MDA level rose (Ercal et al., 2001; Radwan et al., 2010).

The highest level of LPO in soft tissues of the sample collected from El-Balah Road is a high polluted site as compared to the Ring Road in the autumn season (Ghobashy et al, 2021). This may be due to the highest concentration of Cu metal that is found in the autumn season at site II (Ghopashy et al., 2021). This finding is consistent with previous findings that showed Cu exposure caused oxidative stress in snail tissues, as shown by a rise in LPO levels (Valko et al., 2005; Ghobashy et al, 2021). The causes of LPO after Cu exposure are unknown, but we believe that changes in GSH and metallothionein levels may allow free radicals to be released, allowing HO. and O_2_^.-^ radicals to attack double bonds in membrane lipids and cause LPO elevation. Moreover, mitochondrial respiration as the major source of ROS is promoted by LPO and therefore enhances oxidative stress-induced (Ercal et al., 2001).

GSH plays an important role in retaining the cellular redox state and shielding cells from oxidative damage, in addition to being an essential cofactor for GPx and GST activities (Dickinson and Forman, 2002). Since the concentrations of toxic metals (Cu, Zn, and Pb) were low in the study areas (Ghopashy et al., 2021), GSH reached its peak concentration in the summer season in the two locations. These results agree with the findings of Radwan et al. (2010). They found that the GSH level was decreased in the digestive gland of *T. pisana* snails from six different polluted sites in Alexandria city, Egypt. In the previous study, the level of GSH was significantly lower in the snails from the polluted sites than the value obtained from the same species in the reference site by 31.4% (Radwan et al., 2010).

During the autumn, the GSH concentration was reduced significantly at site I for the two species as compared to site II. This could be due to the sudden increase in levels of Fe, Mg, and Ni in site I during the autumn season (Ghobashy et al., 2021). The strong affinity of these elements for the GSH molecule, as stated by Radwan et al. (2010), appears to be a typical response of mollusks to metal exposure, as evidenced by a decrease in GSH content in the digestive gland.

A decrease in GSH content was followed by an increase in LPO levels in the current data. Since GSH functions as a reducing agent and free-radical trapper and is believed to be a cofactor substrate and/or GSH-related enzymes. It is one of the most significant factors in defending against oxidative attacks by ROS such as LPO (Verma et al., 2007). Pb and Cd, both heavy metals, have electron-sharing affinities that can lead to the formation of covalent bonds (Ercal et al., 2001).

At the two sites or through the four seasons, there was no substantial variation in TP levels between the two animals. Unless the total protein level slightly increases at a site I in the winter season, then the concentration of most heavy metals reaches its highest value (Ghopashy et al., 2021). At site II in the spring season, the levels of Cu, Mg, Cd, and B have reached their highest value. These findings contradicted those of Radwan et al. (2008), who observed that giving *E. vermiculata* methiocarb on alternate days for seven days produced a substantial decrease in TP in the tissues.

Furthermore, the current results contradict those of Abdel-Aal (2004), who discovered that when applied to the land snail *E. vermiculata*, niclosamide molluscicidal raised the total proteins more than the control after 24, 48, 72, and 96 hours of care. A discrepancy between the rate of synthesis and the rate of degradation may have contributed to the decrease in TP levels (Gabr et al., 2007).

The toxic effect of chemical compounds may be due to increased energy consumption and/or degradation of cell organelles in handled snails, which could contribute to protein synthesis inhabitation (El Gohary et al., 2011). Proteins are mainly involved in the architecture of the cell. They are also a source of energy during prolonged periods of stress (Radwan et al., 2008). The snails need more energy to detoxify the toxicants and overcome the induced stress while they are stressed. Since snails have a small supply of carbohydrates, proteins are the next best form of nutrition to support the increased demand (Radwan et al., 2008).

The digestive gland of gastropod mollusks is the key organ of metabolism, serving as the primary location for xenobiotic accumulation and biotransformation (Desouky, 2006). In the current data, it was noticed that the histological structure of the gonads and digestive gland of two species of land snails at the two studied sites were more or less similar to each other. The current study’s description of the digestive gland’s histological composition matched that of other researchers (Hamed et al., 2007; Abo-Bakr, 2011; Abd El-Atti et al., 2019). They studied the toxicological and histological changes of the digestive gland of *E. vermiculata*. Moreover, Ali and said (2019)studied the histological change of the digestive gland of *M. obstructa*.

Histological investigations into the digestive gland of two species in the current study confirmed the presence of three main cell types; digestive, calcium, and excretory cells forming the epithelial wall of the digestive tubules. This finding was in harmony with the findings of other authors who observed these kinds of cells in *E. vermiculata* (Abd El-Atti et al., 2020), *H. pomatia* (Chabicovsky et al., 2004), *H. aspersa* (Snyman et al., 2005), *H. vestalis* (Sharaf et al., 2015) and *M. cartusiana* (Abd El-Atti et al., 2019).

However, the styles and associated nomenclature of gastropod digestive gland cells, as well as the functions assigned to them, are a source of debate (Hamed et al., 2007). In the digestive glands of *Littorina littorea*, Zaldibar et al. (2007)identified two cell types: digestive cells and basophilic cells. Beshr (2000), on the other hand, identified digestive, calcium, and thin cells. In the digestive gland of *E. vermiculata*, Heiba et al. (2002) found digestive, excretory, and secretory cells. In the digestive gland of *E. vermiculata*, Hamed et al. (2007)identified four cell types: digestive, calcium, excretory, and thin cells.

These cell types have been given different names by different authors according to their supposed functions (Aioub et al., 2000). They are believed to be primarily responsible for the uptake and digestion of food materials, absorption, and pinocytosis. Other functions that have been indicated for digestive gland cells are participation in shell regeneration and the storage of glycogen and fat. Moreover, the function of the excretory cells involves the synthesis and elaboration of the same digestive enzymes. Also, these cells become excretory only towards the end of their cycle (Aioub et al., 2000).

The current study confirms the absorptive function of *E. vermiculata* digestive cells, which include organelles involved in absorption and intracellular digestion. Also, the digestive cells are numerous in the digestive gland epithelium. The present findings are in agreement with those described by Aioub et al. (2000), who studied the toxicological and histological characteristics of the digestive gland of *E. vermiculata*. Similarly, The apical surface of digestive cells has a distinct microvillus boundary coated with a glycoprotein glycocalyx, and the lateral surfaces of their apical regions are joined together by zonula adherents accompanied by a long and pleated septate junction, according to Hamed et al. (2007). Their cytoplasm granules and the basally-located nuclei of digestive cells in *E. vermiculata* were detected.

In the present study, the digestive cells displayed morphofunctional changes that may be explained by the fact that they were at distinct stages of activity. This follows Hamed et al. (2007), who identified successive phases of absorption, digestion, and fragmentation in the cycle of the digestive cells in the tubules of *E. vermiculata*. Also, Ali et al. (2013)described histological alterations in the digestive gland of *E. vermiculata* as digestive cells were shrunk or completely degenerated.

The excretory cells described in the present study are similar to the same observations that have been reported by Aioub et al. (2000), Hamed et al. (2007), and Ali et al. (2013). According to Lopes et al. (2001), the related reaction of the granules present in *O. atlanticus* digestive and excretory cells shows a near affinity between the two forms of cells. Since excretory cells are formed from digestive cells, this reaction was also observed in other bacteria.

Calcium cells in the current study have the same shape as those described by Hamed et al. (2007). They contain spherules of calcium salts, which have a characteristic birefringence (Lopes et al., 2001). The function of calcium reserves is unknown, it is thought to play a role in several essential calcium-dependent metabolic reactions, including shell creation and repair, pH control in the digestive tract, and reproduction (Lopes et al., 2001).

Heavy metal toxicity can be effectively demonstrated via histopathological investigations, as it determines the target organ of the examined metal. Also, it reveals the impact of exposure to lethal and sublethal levels of contaminants on organ tissues (Abdallah, 2003). The combined metals exhibited a destructive effect on the digestive tubules, where tubular deterioration and disintegration of the digestive cells were observed. Uptake and release effects of contaminants can be seen in histological sections (Abdallah, 2003).

The general architecture of the digestive gland in *E. vermiculata* and *M. obstructa* snails collected from the highly polluted at site I lost its normal appearance when compared to those collected from the less polluted location at site II. Changes in cell-type composition in the digestive gland of slugs and their effect on biomarkers following transplantation between a comparatively unpolluted and a chronically metal-polluted site were discovered by Zaldibar et al. (2008).

Excretory cells have increased in number and size. These findings corroborated those of Chabicovsky et al. (2004), who discovered that heavy metal toxins like Cd, Cu, and Zn cause an increase in the number of digestive gland excretory cells. Similarly, following exposure to oxamyl, Aioub et al. (2000)discovered a rise in the number of excretory cells in the digestive gland of *E. vermiculata*. The current findings were also consistent with those of Hamed et al. (2007), who discovered that when methomyl was applied topically to the land snail *E. vermiculata*, the amount and size of excretory granules increased after 5 and 7 days of therapy.

In the present study, calcium cells decreased in polluted sites and were packed with enlarged calcium spherules, and exhibited pyknotic nuclei (PN). Many calcium cells had their cytoplasm replaced by huge vacuoles containing darkly stained granules. Furthermore, the current findings corroborated those of Aioub et al. (2000), who discovered that after exposure to oxamyl, the number of calcium cells in the digestive gland of *E. vermiculata* decreased significantly. According to Hamed et al. (2007), heavy metal treatment causes a rise in the amount of calcium spherules in the digestive gland of *E. vermiculata*, implying that these cells are adapted for the aggregation and removal of toxic substances.

The normal histological structure of *E. vermiculata* observed in the current study was similar to that described by Abd El-Atti et al (2019; 2020) and Kandil et al. (2020). Histopathological alterations in samples showed that gonad follicles had lost their normal architecture with degeneration of some stages of spermatogenesis and oogenesis. The Acini appeared to be empty and replaced by brown granules, indicating severe disruption of germ cells and more degenerative effects (tissue necrosis). Mohamed et al. (2000), Heiba et al. (2002), and El-Feky et al. (2003) all came to similar conclusions (2009). This research confirms the use of histopathology as a valuable instrument for tracking anthropogenic pollution of terrestrial habitats, since these changes mark a biological endpoint of contaminant exposure.

When these pathological endpoints are assessed in conjunction with the capacity of the animals to accumulate heavy metals in the body tissues (Ghobashy et al., 2021), a clearer picture of the complex interactions between anthropogenic and natural environmental modifiers has emerged. This has ensured early detection of impending environmental problems that may ensue and also averted potential human health tragedies.

In conclusion, the response of *E. vermiculata* and *M. obstructa* to the effect of heavy metals, as shown by bioaccumulation (Ghobashy et al., 2021), biomarkers, and histopathological changes, demonstrates the potential value of this organism in terrestrial pollution biomonitoring studies. The density, morphometric, biochemical, and histology of *Eobania vermiculata* and *Monacha obstructa* were useful as bioindicator at different seasons and can be applicable for ectotoxicological approach of heavy metals. Oxidative stress and the activation of different defenses (enzymatic and non-enzymatic) have received a lot of attention due to different pollution and season. There is consensus about the conservation of different mechanisms involved in the animal’s defenses to be used as biomarkers.

## Competing Interests

The authors declare there are no competing interests. *The funders had no role in study design, data collection and analysis, decision to publish, or preparation of the manuscript*.

## Author contribution statement

All authors contributed equal.

## Notes

### Competing Interest Statement

The authors have declared no competing interest.

